# A Linear Time Solution to the Labeled Robinson-Foulds Distance Problem

**DOI:** 10.1101/2020.09.14.293522

**Authors:** Samuel Briand, Christophe Dessimoz, Nadia El-Mabrouk, Yannis Nevers

**Affiliations:** Département d’informatique et de recherche opérationnelle (DIRO), Université de Montréal; Department of Computational Biology, University of Lausanne; Center for Integrative Genomics, University of Lausanne; Centre for Life’s Origins and Evolution, Genetics Evolution and Environment, University College London; Department of Computer Science, University College London; SIB Swiss Institute of Bioinformatics

**Keywords:** Tree Metric, Gene Trees, Labeled Trees, Robinson-Foulds, Edit Distance

## Abstract

**Motivation:** Comparing trees is a basic task for many purposes, and especially in phylogeny where different tree reconstruction tools may lead to different trees, likely representing contradictory evolutionary information. While a large variety of pairwise measures of similarity or dissimilarity have been developed for comparing trees with no information on internal nodes, very few address the case of inner node-labeled trees. Yet such trees are common; for instance reconciled gene trees have inner nodes labeled with the type of event giving rise to them, typically speciation or duplication. Recently, we proposed a formulation of the Labeled Robinson Foulds edit distance with edge extensions, edge contractions between identically labeled nodes, and node label flips. However, this distance proved difficult to compute, in particular because shortest edit paths can require contracting “good” edges, i.e. edges present in the two trees.

**Results:** Here, we report on a different formulation of the Labeled Robinson Foulds edit distance — based on node insertion, deletion and label substitution — which we show can be computed in linear time. The new formulation also maintains other desirable properties: being a metric, reducing to Robinson Foulds for unlabeled trees and maintaining an intuitive interpretation. The new distance is computable for an arbitrary number of label types, thus making it useful for applications involving not only speciations and duplications, but also horizontal gene transfers and further events associated with the internal nodes of the tree. To illustrate the utility of the new distance, we use it to study the impact of taxon sampling on labeled gene tree inference, and conclude that denser taxon sampling yields better trees.

## 1 Introduction

Gene trees are extensively used, not only for inferring phylogenetic relationships between corresponding taxa, but also for inferring the most plausible scenario of evolutionary events leading to the observed gene family from a single ancestral gene copy. This has important implications towards elucidating the functional relationship between gene copies. For this purpose, reconciliation methods (reviewed in Boussau and Scornavacca, 2020) embed a given gene tree into a known species tree. This process results in the labeling of the internal nodes of the gene tree with the type of events which gave rise to them, typically speciations and duplications, but also horizontal gene transfers or possibly other events (whole genome duplication, gene convergence, etc). For example, information on duplication and speciation node labeling is provided for the trees of the Ensembl Compara database (Vilella et al., 2009).

The existence of a variety of different phylogenetic inference methods leading to different, potentially inconsistent, trees for the same dataset, brings forward the need for appropriate tools for comparing them. Although comparing labeled gene trees remains a largely unexplored field, a large variety of pairwise measures of similarity or dissimilarity have been developed for comparing unlabeled evolutionary trees. Among them are the methods based on counting the structural differences between the two trees in terms of path size, bipartitions or quartets for unrooted trees, clades or triplets for rooted trees (Cardona et al., 2010; Critchlow et al., 1996; Estabrook et al., 1985), or those based on minimizing a number of rearrangements that disconnect and reconnect subpieces of a tree, such as nearest neighbour interchange (NNI), subtree-pruning-regrafting (SPR) or Tree-Bisection-Reconnection (TBR) moves (BL Allen and Steel, 2001; Hickey et al., 2008; Jiang et al., 2000). While the latter methods are NP-hard (Lin et al., 2012), the former are typically computable in polynomial time. In particular, the Robinson-Foulds (*RF*) distance, defined in terms of bipartition dissimilarity for unrooted trees, and clade dissimilarity for rooted trees (Mittal and Munjal, 2015), can be computed in linear (Day, 1985), and even sublinear time (Pattengale et al., 2007).

On the other hand, metrics have also been developed for node-labeled trees (rooted, and sometimes with an order on nodes) arising from many different applications in various fields (parsing, RNA structure comparison, computer vision, genealogical studies, etc), where node labels in a given tree are pairwise different i.e. no repeated labels). For such trees, the standard Tree Edit Distance (TED) (Zhang and Shasha, 1989), defined in terms of a minimum cost path of node deletion, node insertion and node change (label substitution) transforming one tree to another, has been widely used. While the general version of the problem on unordered labeled trees with a non-constant cost function on edit operations is NP-complete (Zhang, Statman, et al., 1992), most variants are solvable in polynomial time (Schwarz et al., 2017; Zhang, 1993, 1996).

The metric we developed in Briand, Dessimoz, El-Mabrouk, Lafond, et al., 2020, referred to as *ELRF*, is the first effort towards comparing labeled gene trees, expressed in terms of trees with a binary node labeling (typically speciation and duplication). *ELRF* is an extension of the *RF* distance, one of the most widely used tree distance, not only in phylogenetics, but also in other fields such as in linguistics, for its computational efficiency, intuitive interpretation and the fact that it is a true metric. Improved versions of the *RF* distance have also been developed (Lin et al., 2012; Moon and Eulenstein, 2018) to address the distance’s drawbacks, which are lack of robustness (a small change in a tree may cause a disproportional change in the distance) and skewed distribution. Classically defined in terms of bipartition or clade dissimilarity, the *RF* distance can similarly be defined in terms of edit operations on tree edges: the minimum number of edge contraction and extension needed to transform one tree into the other (Robinson and Foulds, 1981). In Briand, Dessimoz, El-Mabrouk, Lafond, et al., 2020, this definition of the *RF* distance was extended to trees with binary node labeling by including a node *flip* operation, alongside edge contractions and extensions. While remaining a metric, *ELRF* turned out to be much more challenging to compute. As a result, we proposed a heuristic to compute it efficiently.

In this paper, we explore a different extension of *RF* to node-labeled trees with labels belonging to a set of label types, directly derived from TED (Zhang and Shasha, 1989), which is a reformulation of the *RF* distance in terms of edit operations on tree nodes rather than on tree edges. We show that this new distance is computable in linear time for an arbitrary number of label types, thus making it useful for applications involving not only speciations and duplications, but also horizontal gene transfers and further events associated with the internal nodes of the tree. We show that the new distance compares favourably to *RF* and *ELRF* by performing simulations on labeled gene trees of 182 leaves. Finally, we use our new distance in the purpose of measuring the impact of taxon sampling on labeled gene tree inference, and conclude that denser taxon sampling yields better predictions.

## 2 Notation and Concepts

The Robinson-Foulds (RF) distance is defined in the literature for rooted and unrooted trees. Moreover, as mentioned in Briand, Dessimoz, El-Mabrouk, Lafond, et al., 2020, the problem of computing the RF distance for two rooted trees can be reduced to computing the RF distance for the two corresponding unrooted trees obtained by grafting an edge linking the root to a dummy leaf. Therefore, in this paper we restrict ourselves to unrooted trees. We begin by introducing few required notations.

Let *T* be a tree with node set *V* (*T*) and edge set *E*(*T*). Given a node *x* of *T*, the *degree of x* is the number of edges incident to *x*. In this paper, the considered trees are unrooted with all internal nodes being of degree at least 3. An internal node of degree 3 is said to be *binary*.

We denote by *L*(*T*) *⊆ V* (*T*) the set of *leaves of T*, i.e. the set of nodes of *T* of degree one. Given a set *ℒ* (let us say taxa or genetic elements), a tree *T* on *ℒ* is a tree with a one-to-one relationship between *L*(*T*) and *ℒ*.

A node of *V* (*T*) \ *L*(*T*) is called an *internal node*. A tree with a single internal node *x* is called a *star tree*, and *x* is called a *star node*. An edge connecting two internal nodes is called an *internal edge*; otherwise, it is a *terminal edge*. Moreover, a *rooted tree* admits a single internal node *r*(*T*) considered as the root.

We call *N* (*x*) = *{y* : *{x, y} ∈ E*(*T*)*}* the set of neighbours of an internal node *x* of *T*.

A *subtree S* of *T* is a tree such that *V* (*S*) *⊆ V* (*T*), *E*(*S*) *⊆ E*(*T*) and any edge of *E*(*S*) connects two nodes of *V* (*S*).

The *bipartition* of an unrooted tree *T* corresponding to an edge *e* = *{x, y}* is the unordered pair of clades *L*(*T*_*x*_) and *L*(*T*_*y*_) where *T*_*x*_ and *T*_*y*_ are the two subtrees rooted respectively at *x* and *y* obtained by removing *e* from *T*. We denote by *ℬ* (*T*) the set of non-trivial bipartitions of *T*, i.e. those corresponding to internal edges of *T*.

### 2.1 The Robinson-Foulds distance

Given two unrooted trees *T* and *T ′* on *ℒ*, the Robinson-Foulds (*RF*) distance between *T* and *T*^*′*^is the size of the symmetric difference between the bipartitions of the two trees. More precisely,

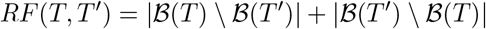

The *RF* distance is equivalently defined in terms of an edit distance on edges. However, as for node-labeled trees an additional substitution operation on node labels will be required, for the sake of standardization, we reformulate the edit operations to operate on nodes rather than on edges.

#### Definition 1

(node edit operations). *Two edit operations on the nodes of a tree T are defined as follows:*

- ***Node deletion (****Del*): *Let x be an internal node of T which is not a star node and y be a neighbour of x which is not a leaf. Deleting x with respect to y means making the neighbours of x become neighbours of y. More precisely, Del*(*T, x, y*) *is an operation transforming the tree T into the tree T ′ obtained from T by removing the edge {x, z} for each z ∈ N* (*x*), *creating the edge {y, z} for each z ∈ N* (*x*) | *{y}, and then removing node x*.
- ***Node insertion (****Ins*): *Let y be an internal node of V* (*T*) *of degree at least 3. Inserting x as a neighbour of y entails making x the neighbour of a subset Z* ⊊ *N* (*y*) *such that* |*Z*| *≥* 2. *More precisely, Ins*(*T, x, y, Z*) *is an operation transforming the tree T into the tree T obtained from T by removing the edges {y, z*_*i*_*}, for all z*_*i*_ *∈ Z, creating a node x and a new edge e* = *{x, y}, and creating new edges {x, z*_*i*_*}, for all z*_*i*_ *∈ Z*.

Formally, let *T* and *T ′* be two trees on the same *ℒ*. The *Robinson-Foulds* or *Edit distance* (Robinson and Foulds, 1981) *RF* (*T, T ′*) between *T* and *T ′* is the size of a shortest path of edge edit operations (i.e. *edge extensions and edge contractions)* transforming *T* into *T*. This distance measure, equivalently defined as the size of the symmetric difference between the non-trivial bipartitions of the two trees, has been shown to be a metric. Notice the one-to-one correspondence between operations on nodes and operations on edges. In fact, deleting a node *x* by an operation *Del*(*T, x, y*) results in removing the edge *{x, y}*, while inserting a node *x* by an operation *Ins*(*T, x, y, Z*) results in inserting the edge *{x, y}*. Here, we define the *RF* distance in terms of edit operations on nodes.

Call a *bad edge* of *T* with respect to *T ′* (or similarly of *T*^*′*^with respect to *T*; if there is no ambiguity, we will omit the “with respect to” precision) an edge representing bipartitions which are not shared by the two trees, i.e. an edge of *T* (respec. *T*^*′*^) defining a bipartition of *ℬ* (*T*) (respec. *ℬ* (*T* ^*′*^)) which is not in *ℬ* (*T ′*) (respec. in *ℬ* (*T*)). An edge which is not bad is said to be *good*. Terminal edges are always good.

## 3 Generalizing the Robinson-Foulds distance to Labeled Trees

A tree *T* is *labeled* if and only if each internal node *x* of *T* has a label *λ*(*x*) *∈* Λ, Λ being a finite set of labels. For gene trees, labels usually represent the type of event leading to the bifurcation, typically duplications and speciations, although other events, such as horizontal gene transfers, may be considered. The metric defined in this paper holds for an arbitrary number of labels. We generalize the *RF* distance to labeled trees by generalizing the edit operations defined above. This is simply done by introducing a third operation for node labels editing.

### Definition 2

(Labeled node edit operations). *Three edit operations on internal nodes of a labeled tree T are defined as follows:*

- ***Node deletion:*** *Del*(*T, x, y*) *is an operation deleting an internal node x of T with respect to a neighbour y of x which is not a leaf, defined as in Definition 1*.
- ***Node insertion:*** *Ins*(*T, x, y, Z, λ*) *is an operation inserting an internal node x as a new neighbour of a non-binary node y, and moving Z* ⊊ *N* (*y*) *such that* |*Z*| *≥* 2, *to be the neighbours of x, as defined in Definition 1. In addition, the inserted node x receives a label λ ∈* Λ.
- ***Node label substitution:*** *Sub*(*T, x, λ*) *is an operation substituting the label of the internal node x of T with λ ∈* Λ.

These operations are illustrated in Figure 1.

**Figure 1.**
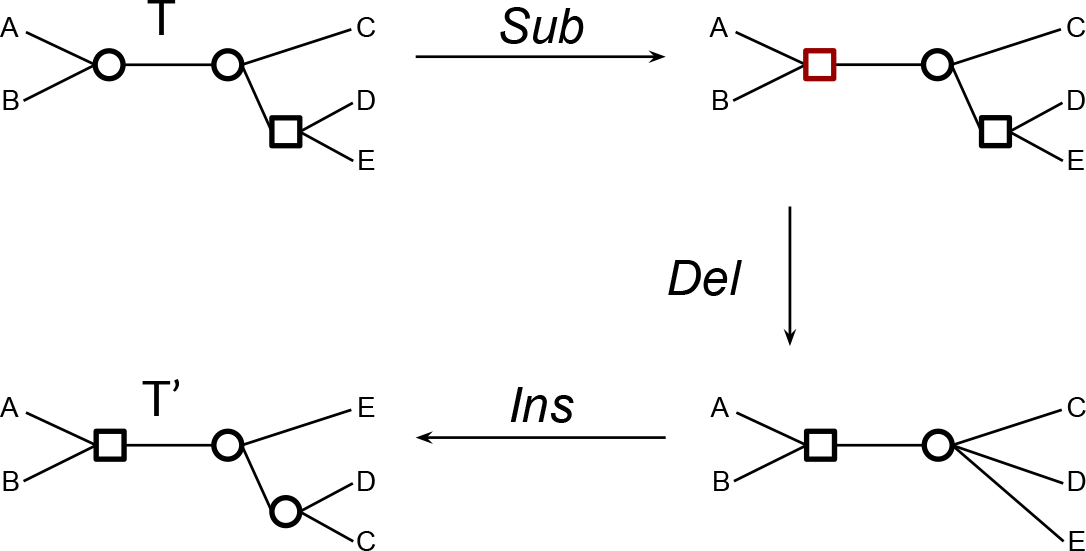
The transformation of a tree *T* into a tree *T ′* depicting the three edit operations on nodes. From top to bottom: node label substitution (leading to the red label), node deletion (the parent of *D* and *E*) and node insertion (the parent of *D* and *C*).

Let *T* _*ℒ*_ be the set of unrooted and labeled trees on *ℒ*. For two trees *T, T ′* of *T* _*ℒ*_, we call the *Labeled Robinson Foulds* distance between *T* and *T* and denote by *LRF* (*T, T* ^*′*^) the size of a shortest path of labeled node edit operations transforming *T* into *T ′* (or vice versa). The two following lemmas state that, similarly to *RF, LRF* is a true metric. Moreover, *LRF* is exactly *RF* for unlabeled trees (or similarly labeled with a single label).

In the following, the *unlabeled version of* a tree *T ∈ 𝒯*_*ℒ*_ is simply *T* ignoring its node labels.

### Lemma 1.

*The function LRF* (*T, T*^*′*^) *assigning to each pair* 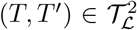 *the size of a shortest path of node edit operations transforming T into T ′ defines a distance on T*_*L*_.

*Proof*. The non-negative, identity and triangular inequality conditions are obvious. For the symmetric condition, notice that we can reverse every edit operation in a path from *T ′* to *T* to obtain a path from *T* to *T ′* with the same number of events, and vice versa (insertions and deletions are symmetrical operations, and any substitution can be reversed by a substitution). We thus have *LRF* (*T*^*′*^, *T*) *≤ LRF* (*T, T ′*) and *LRF* (*T, T*) *≤ LRF* (*T ′, T*), and equality follows.

The next lemma directly follows from the fact that node substitutions are never applied in case of a label set restricted to a single label.

### Lemma 2.

*If* Λ *is restricted to a single label, then for each pair* 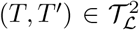,*LRF*(*T, T*′) = *RF*(*T, T*′).

A previous extension of *RF* to labeled trees, based on edit operations on edges rather than on nodes, was introduced in Briand, Dessimoz, El-Mabrouk, Lafond, et al., 2020. This distance, which we call *ELRF*, was defined on three operations:

- Edge extension *Ext*(*T, x, X*) creating an edge *{x, y}* and defined as a node insertion *Ins*(*T, y, x, X, λ*(*x*)) inserting a node *y* as a neighbour of *x* and assigning to *y* the label of *x*;
- Edge contraction *Cont*(*T, {x, y}*) is equal to a node deletion *Del*(*T, y, x*) deleting *y*, but contrary to LRF, requires that *λ*(*x*) = *λ*(*y*);
- Node flip *Flip*(*T, x, λ*) assigning the label *λ* to *x*.

Given two labeled trees *T* and *T ′* of *T* _*ℒ*_, *ELRF* (*T, T ′*) is the size of the shortest path of edge extension, edge contraction and label flip required to transform *T* to *T ′*.

The following lemma makes the link between *LRF* and *ELRF*.

### Lemma 3.

*For any pair* 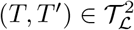,

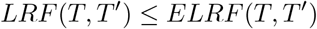

*Proof*. Let *𝒫* be a path of edge edit operations and label flip transforming *T* into *T ′* such that | *𝒫*| = *ELRF* (*T, T ′*). Then the sequence *𝒫* obtained from *𝒫 ′* by replacing each edge extension by the corresponding node insertion, each edge contraction by the corresponding node deletion and each node flip by the corresponding node substitution is clearly a path of node edit operations of size | *𝒫* ^*′*^ | = | *𝒫′* | = *ELRF* (*T, T′*) transforming *T* into *T ′*. And thus *LRF* (*T, T ′*) *≤ ELRF* (*T, T*^*′*^). Figure 2 depicts an example were the inequality is strict.

**Figure 2.**
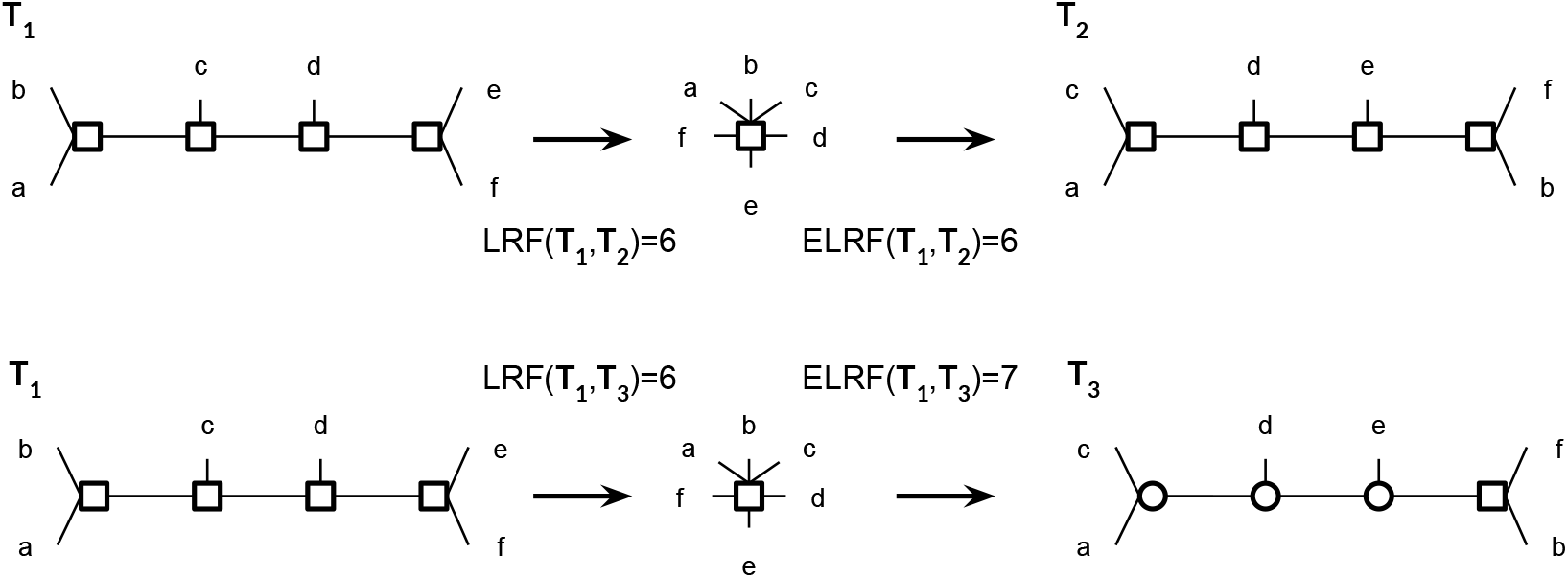
The transformation of a tree *T*_1_ into a tree *T*_2_ (respect. *T*_3_) depicting where the equality (respect. the inequality) is strict between *LRF* (*T*_1_, *T*_2_) and *ELRF* (*T*_1_, *T*_2_) (respect. *LRF* (*T*_1_, *T*_3_) and *ELRF* (*T*_1_, *T*_3_)).

The rest of this paper is dedicated to computing the edit distance *LRF* (*T, T*^*′*^) for any pair (*T, T*^*′*^) of trees of *𝒯* _*ℒ*_.

### 3.1 Reduction to Islands

In this section, we define a subdivision of the two trees into pairs of maximal subtrees that can be treated separately.

While a good edge *e* in *T* has a corresponding good edge *e* in *T ′* (the one defining the same bipartition), a bad edge in *T* has no corresponding edge in *T ′*. However, these bad edges may be grouped into pairs of corresponding *islands* (called maximum bad subtrees in Briand, Dessimoz, El-Mabrouk, Lafond, et al., 2020), as defined bellow.

#### Definition 3

(Islands). *An* island *of T is a maximal subtree I such that all its internal edges are bad edges of T, and all terminal edges of I are good edges of T*. *The* size *of I, denoted ϵ* (*I*), *is its number of internal edges*.

In other words, an island of *T* is a maximal subtree with all internal edges (if any) being bad edges of *T*, and all terminal edges being good edges of *T*. Notice that an island *I* of *T* may have no internal edge at all, i.e. it may be restricted to a star tree (if *ϵ* (*I*) = 0). Notice also that each bad edge of *T* belongs to a single island, while each good edge belongs to exactly two islands of *T* if it is an internal edge of *T*, or to a single island if it is a terminal edge of *T*.

Finally, the following lemma (lemma 3 from Briand, Dessimoz, El-Mabrouk, Lafond, et al., 2020) shows that there is a one-to-one correspondence between the islands of *T* and those of *T*^*′*^.

#### Lemma 4.

*Let I be an island of T with the set {e*_*i*_*}*_1*≤i≤k*_ *of terminal edges, and let* 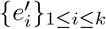 *be the corresponding set of edges in T ′*. *Then the subtree I ′ of T*^*′*^, *containing all ei edges as terminal edges, is unique. Moreover, it is an island of T*^*′*^.

*Proof*. As 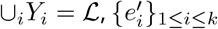 are the only terminal edges of any subtree *I ′* of *T ′* containing the set 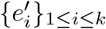 as terminal edges. As *T* is a tree, for any 1 *≤ i* ≠ *j ≤ k*, there is only one possible path from 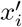 to 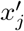. Uniqueness follows.

Suppose that such a subtree *I ′* is not an island. Then it contains an internal good edge *e ′* = (*x* ^*′*^, *y* ^*′*^). In other words, there is a non-trivial bipartition of *{Y*_*i*_*}*_1*≤i≤k*_ which is also a bipartition in *I*. This contradicts the fact that *I* is an island of *T*. Finally, as all terminal edges of *I* are good edges of *T*^*′*^, it follows that *I ′* is an island of *T′*.

For any island *I* of *T*, let *I ′* be the corresponding island of *T′*. We call (*I, I ′*) an *island pair* of; (*T, T* ^*′*^). See Figure 3 for an example.

**Figure 3.**
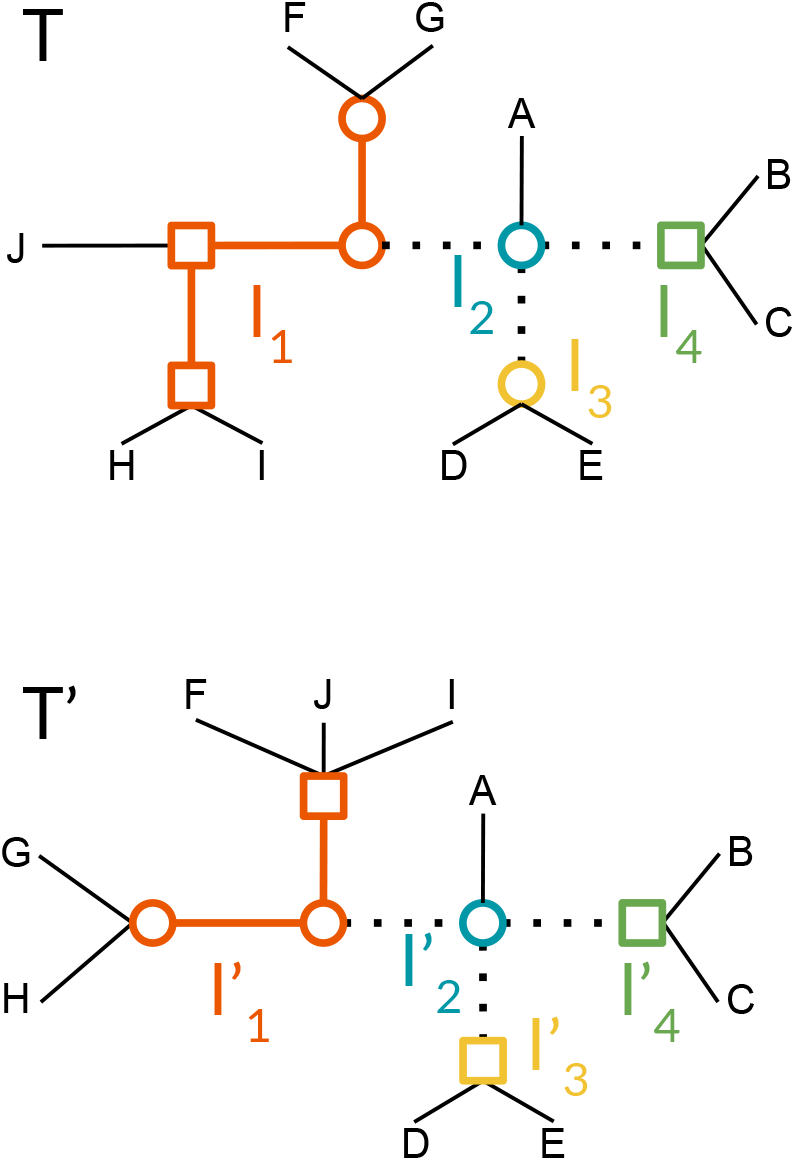
Two trees *T* and *T* ^*′*^on *𝒯 ℒ* for ℒ = *A, B, C, D, E, F, I, J*, with a binary labeling of internal nodes (squares and circles). Dotted lines represent good internal edges, solid lines represent bad edges and thin lines represent terminal edges (which are good edges). This representation highlights the subdivision of the two trees into the island pairs 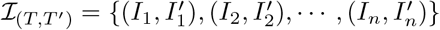. Notice that each dotted line is a terminal edge of its two adjacent islands.

Now, let 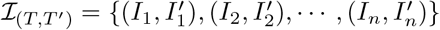 be the set of island pairs of (*T, T ′*). For 1 *≤ i ≤ n*, let *𝒫*_*i*_ be a shortest path of labeled node edit operations transforming *I*_*i*_ into 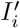. Then the path *𝒫* obtained by performing consecutively *𝒫*_1_, *𝒫*_2_, …, *𝒫*_*n*_ (that we represent later as *𝒫*_1_. *𝒫*_2_. …. *𝒫*_*n*_) clearly transforms *T* into *T ′*. Therefore we have

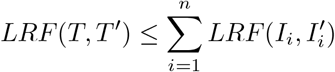

As described in Briand, Dessimoz, El-Mabrouk, Lafond, et al., 2020, one major issue with ELRF is that good edge contractions may not be avoided in a shortest path of edit operations transforming *T* into *T*, resulting in island merging. In other words, treating island pairs separately may not result in an optimal scenario of edit operations under *ELRF*, preventing the above inequality from being an equality. Interestingly, the equality holds for the *LRF* distance, as we show in the next section.

### 3.2 Computing the *LRF* distance on islands

We require an additional definition. Two trees *I* and *I ′* of an island pair are said to *share a common label l ∈* Λ if there exist *x ∈ V* (*I*) and *x ′ ∈ V* (*I ′*) such that *λ*(*x*) = *λ*(*x′*) = *l*. If *I* and *I* do not share any common label, then (*I, I′*) is called a *label-disjoint* island pair. For example, the pair 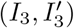 in Figure 3 or the pair (*I, I ′*) in Figure 4 are label-disjoint.

**Figure 4.**
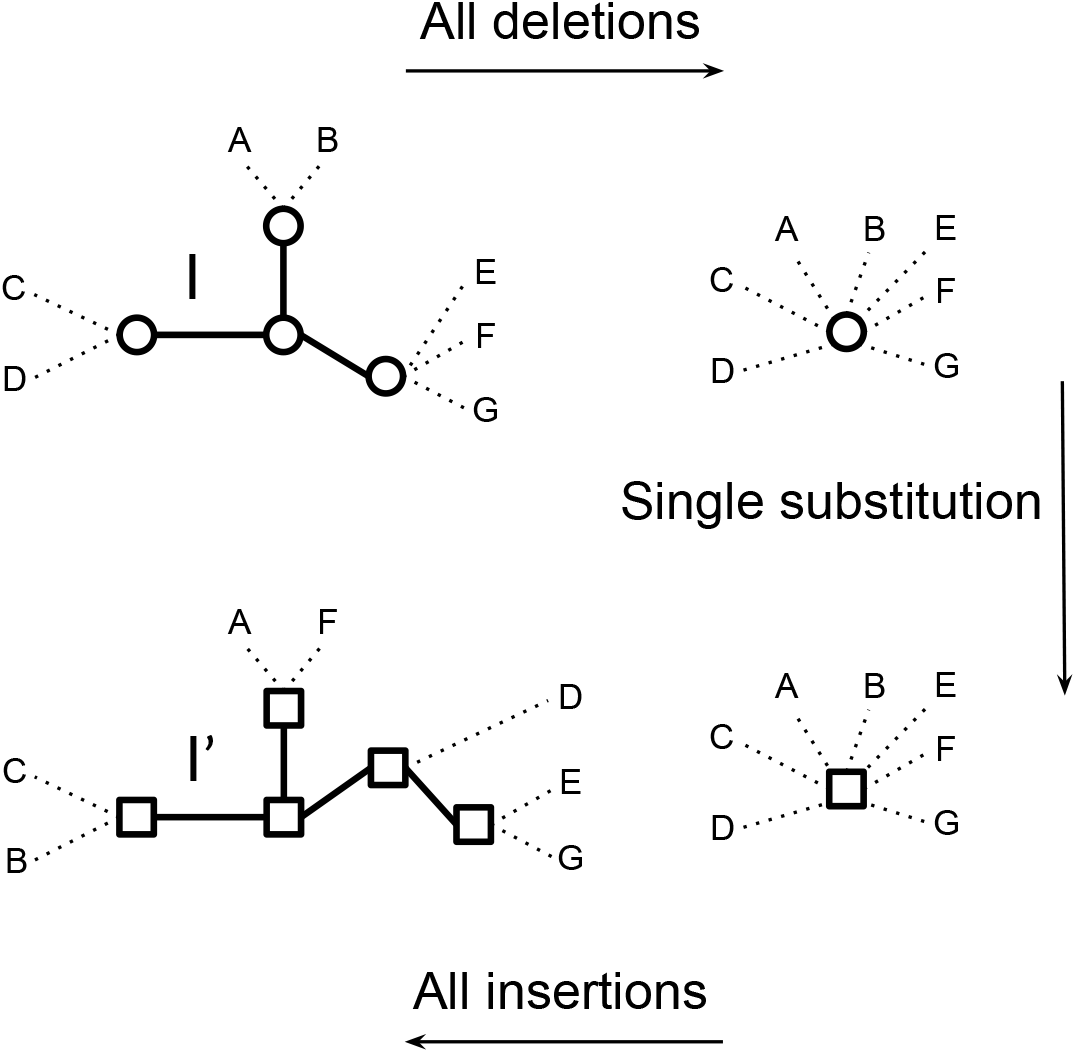
An optimal sequence of edit operations for the island pair (*I, I* ^*′*^).

Now let (*I, I*^*′*^) be an island pair. Transforming *I* into *I ′* can be done by reducing *I* into a star tree by performing a sequence of node deletions (if any, i.e. if *I* is not already a star tree), and then raising the star tree by inserting the required nodes to reach *I ′*. Only the unique node not deleted during the first step might require a label substitution; for all inserted nodes, the label can be chosen to match that of *I′*. However, if *I* and *I ′* share a common label *l* among their internal nodes, then the deletions can be done in a way such that the surviving node *x* of *I* is one with label *λ*(*x*) = *l*, thus avoiding the need for any substitution. The number of required operations is thus *ϵ* (*I*) deletions, followed by zero or one substitution, followed by *ϵ* (*I* ^*′*^) insertions. Alternatively, the problem can be seen as one of reducing the two trees into star trees by performing *ϵ* (*I*)+ *ϵ* (*I* ^*′*^) deletions, in a way reducing the two islands into two star trees sharing the same label, if possible. Figure 4 depicts an example of such tree editing for a label-disjoint island pair.

The following lemmas show that the sequential way of doing described above is optimal.

#### Lemma 5.

*Let* (*I, I*^*′*^) *be an element of I*_(*T,T′*,)_. *Then:*

- *If I and I ′ share a common label, then LRF* (*I, I*^*′*^) = *ϵ* (*I*) + *ϵ* (*I*).
- *Otherwise LRF* (*I, I*^*′*^) = *ϵ* (*I*) + *ϵ* (*I ′*) + 1.

*Proof*. The scenario depicted above for transforming *I* into *I* clearly requires *ϵ* (*I*) + *ϵ* (*I ′*) node insertions and deletions, and an additional node label substitution in case *I* ans *I* are label-disjoint. We can conclude that *LRF* (*I, I ′*) *≤ϵ* (*I*) + *ϵ* (*I ′*) if *I* and *I*^*′*^ share a common label and *LRF* (*I, I ′*) *≤ϵ* (*I*) + *ϵ* (*I ′*) + 1, if *I* and *I* are label-disjoint.

On the other hand, as all the edges of *I* are bad edges, they should be all removed, before reinserting those of *I*. Now, since an edit operation can remove or insert at most one edge, and the only operations removing an edge are node removal or node insertion, we clearly require at least *ϵ* (*I*)+ *ϵ* (*I*^*′*^) node removals and insertions to transform the unlabeled form of the tree *I* into the unlabeled form of *I*. Furthermore, as deletions do not affect star nodes, at least one node in *I* should survive (i.e. not be affected by a node deletion). Thus, if the two trees are label-disjoint, then at least one node label substitution is required. We can then conclude that *LRF* (*I, I ′*) *≥ϵ* (*I*) + *ϵ* (*I ′*) if *I* and *I ′* share a common label and *LRF* (*I, I*^*′*^) *≥ϵ* (*I*) + *ϵ* (*I ′*) + 1, if *I* and *I* are label-disjoint, which concludes the proof.

We have obviously 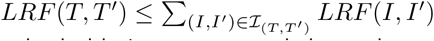 It remains to show that the symmetrical inequality also holds, i.e. we cannot do better by merging islands, and thus pairs of islands can be considered separately. The following lemma states that we can always find a sequence of operations, at each step maintaining or increasing the number of islands, i.e. never merging islands.

For a path *P* = (*o*_1_, *o*_2_, *…o*_*p*_) transforming a tree *T* into a tree *T ′* and 1 *≤ k ≤ p*, denote by *T*_*k*_ the tree obtained from *T* after performing the sub-sequence of operations *𝒫* _*k*_ = (*o*_1_, *… o*_*k*_).

#### Lemma 6.

*Let T and T be two trees of T* _*ℒ*_. *There is a shortest path 𝒫* = (*o*_1_, *o*_2_, *…o*_*p*_) *of edit operations transforming T into T such that for each k*, 2 *≤ k ≤ p*, |*I*(*T*_*k−*1_, *T ′*)| *≤* | *ℒ* (*T*_*k*_, *T ′*)|.

*Proof*. Let *P* = (*o*_1_, *o*_2_, *…o*_*p*_) be a shortest path transforming *T* into *T ′*, Denote 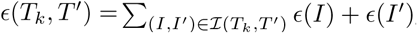 vand *ξ*(*T*_*k*_, *T*) the number of label-disjoints pairs of *ℒ* (*T*_*k*_, *T ′*). Assume *P* contains an operation reducing the number of islands of *T*_*i−*1_, and let *o*_*i*_ be the last operation of that form, i.e. |*I*(*T*_*i−*1_, *T ℒ*)| *>* | *ℒ* (*T*_*i*_, *T ′*)|. Such an operation can only be a deletion *Del*(*T*_*i−*1_, *x, y*) where *e* = *{x, y}* is a good edge, thus merging the two islands *I*_*x*_, *I*_*y*_ containing this good edge.

As, by assumption, *o*_*i*_ is the last operation merging two islands, at that point each pair of islands is treated separately, and we deduce from the fact that 𝒫is a shortest path that *LRF* (*T*_*i*_, *T*) = Σ (*I,I ′*,) *∈I*(*Ti,T*,) *LRF* (*I, I*), and thus

*LRF* (*T*_*i−*1_, *T*) = 1 + 1 + *ϵ* (*T*_*i*_, *T ′*) + *ξ*(*T*_*i*_, *T ′*).

On the other hand, there is a path from *T*_*i−*1_ to *T ′*of size *c*(*T*_*i−*1_, *T*) = *ϵ* (*T*_*i−*1_, *T ′*) + (*I,I*,)*∈I*(*Ti,T*,) *LRF* (*I, I*). Then, from Lemma 5, *LRF* (*Ti−*1, *T ′*) = *ξ*(*T*_*i−*1_, *T*^*′*^).

As *o*_*i*_ is a deletion of a good edge *e* = *{x, y}*, it destroys the bipartition defined by this edge in *T*_*i−*1_, consequently the corresponding edge in *T* becomes a bad edge. Therefore *E*(*T*_*i−*1_, *T*) = *E*(*T*_*i*_, *T*) *−* 1.

On the other hand, let *δ* = *ξ*(*T*_*i*_, *T*)*−ξ*(*T*_*i−*1_, *T*^*′*^) be the difference between the number of label-disjoint pairs of islands after performing the operation *o*_*i*_ merging two pairs of islands 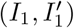 and 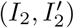

- If both pairs (*I*_1_, *I*1) and (*I*_2_, *I*2) share a common label, then the merged pair also shares a common label, and thus *Δ* = 0;
- If both pairs 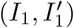 and 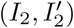 are label-disjoint, then after the merging the resulting pair of islands may or may not share a common label and thus *−*2 *≤ Δ ≤ −*1;
- If only one of the two pairs 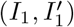 and 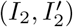 share a common label, then after the merging, the resulting pair of islands may or may not share a common label and thus *−*1 *≤ Δ ≤* 0;

Therefore, in all cases we have *ξ*(*T*_*i*_, *T* ^*′*^) *≤ ξ*(*T*_*i−*1_, *T ′*) *≤ ξ*(*T*_*i*_, *T ′*) + 2.

Recall *c*(*T*_*i−*1_, *T*) = *ϵ* (*T*_*i−*1_, *T ′*) + *ξ*(*T*_*i−*1_, *T ′*) = *ϵ* (*T*_*i*_, *T ′*) *−* 1 + *ξ*(*T*_*i−*1_, *T ′*).

Thus *c*(*T*_*i−*1_, *T ′*) *≤ϵ* (*T*_*i*_, *T ′*) *−* 1 + *ξ*(*T*_*i*_, *T ′*) + 2 = *ϵ* (*T*_*i*_, *T ′*) + *ξ*(*T*_*i*_, *T ′*) + 1 = *LRF* (*T*_*i−*1_, *T ′*). As *LRF* (*T*_*i−*1_, *T ′*) is the size of the shortest path from *T*_*i−*1_ to *T ′*, we should have *c*(*T*_*i−*1_, *T ′*) = *LRF* (*T*_*i−*1_, *T*^*′*^).

Therefore, replacing the sequence of operations (*o*_*i*_, *…o*_*p*_) on *T*_*i−*1_ by a sequence of operations solving each pair if islands separately leads to the same number of operations.

We are now ready to prove the equality leading to the efficient computation of the *LRF* distance of two trees (see Figure 5 for an example).

**Figure 5.**
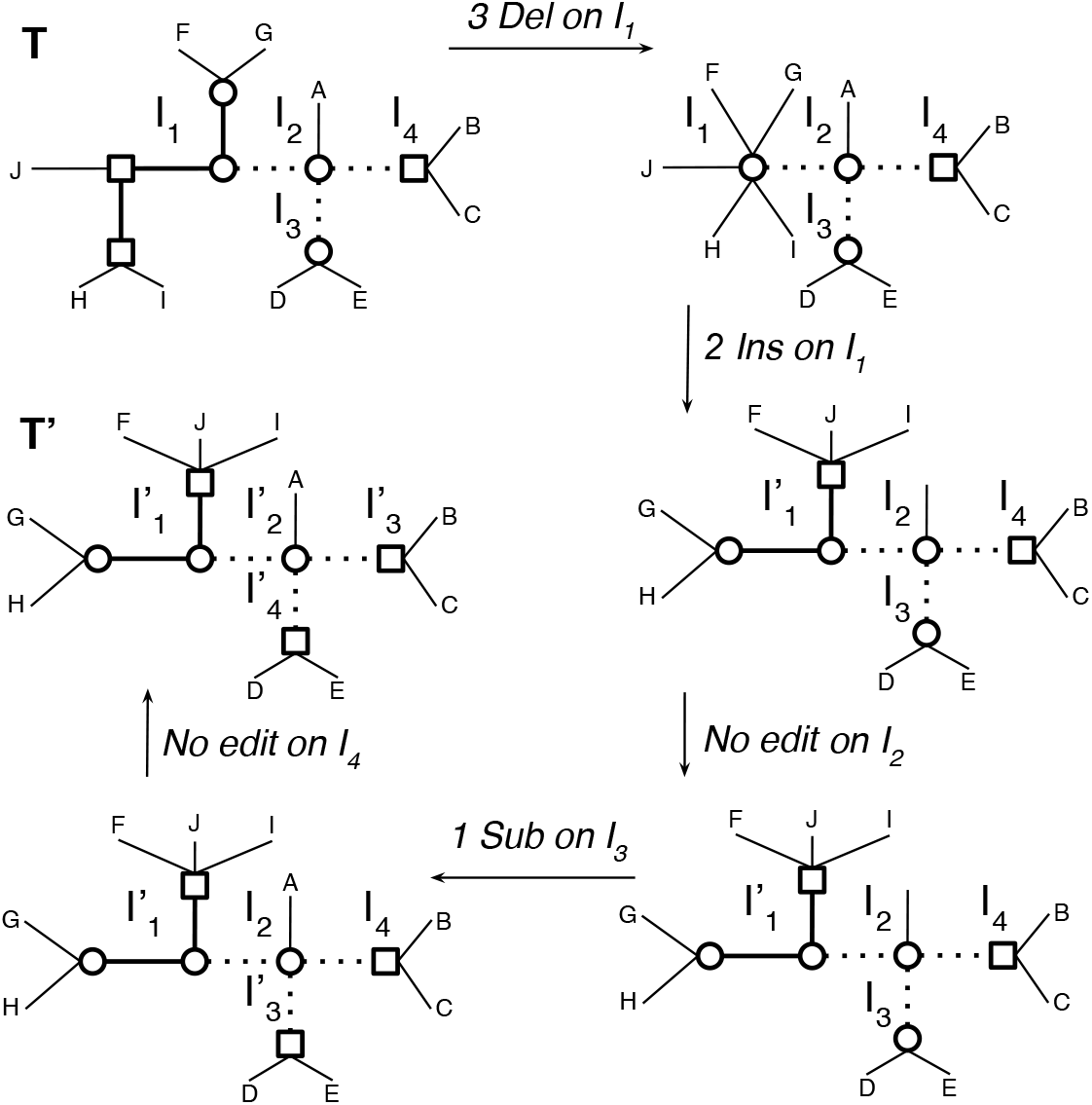
A path *P* transforming *T* into *T ′*of the form *𝓅*_1_. *𝓅*_2_. *𝓅*_3_. *𝓅*_4_, each *𝓅*_*i*_ being a shortest path for the island pair 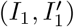. Here | *𝓅*_1_| = 6, | *𝓅*_2_| = 0, | *𝓅*_3_| = 1, and | *𝓅*_4_| = 0.

#### Theorem 1.

*Let* 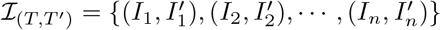 *be the island pairs of T and T ′*. *Then*

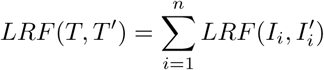

*Proof*. Let *𝓅* be a shortest path transforming *T* into *T ′* verifying the condition of Lemma 6, i.e. not involving any removal of good edges. As islands can only share good edges, and good edges are never removed by any operation of *P*, islands are never merged during the process of transforming *T* into *T ′*, and thus can be treated separately. Let *𝓅*_*i*_, 1 *≤ i ≤ n*, be the subpath of edit operations transforming *I*_*i*_ into 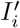 Each *𝓅*_*i*_ should be a shortest path from *I*_*i*_ to 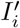as otherwise it can be replaced by a shortest path, contradicting the fact that *P* is a shortest path.

The next result directly follows from Lemma 5 and Theorem 1.

#### Corollary 1.

*Let* 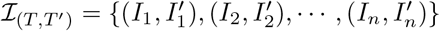 *be the island pairs of T and T ′ and Δ be the number of label-disjoint pairs. Then*

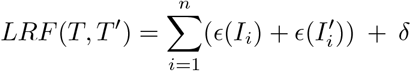

## 4 Algorithm

We present our algorithm for computing the *LRF* distance (Algorithm 1). The input is a pair of trees *T*_1_, *T*_2_ of *T*_*L*_. We show that *LRF* (*T*_1_, *T*_2_) can be computed in time *𝒪* (*n*), where *n* = | *ℒ*|.

### 4.1 The *LRF* () function

We start with the identification of good edges. Lines 1 and 2 of Algorithm 1 retrieve the non-trivial bipartitions for each input tree and Line 3 intersects the obtained bipartitions of *T*_1_ and *T*_2_ to generate the set of good edges shared by the two input trees. This is the same procedure as to compute the conventional (unlabeled) Robinson-Foulds distance, so we do not detail it here. It is sufficient to know that it can be done in time *𝒪* (*n*) (Day, 1985).

Next the algorithm identifies and characterises the islands of *T*_1_ and *T*_2_ (lines 4 and 5). This is performed using an auxiliary function *getIslands*() (Algorithm 2), which we describe in detail below. As we shall see, it runs in *𝒪* (*n*).

Next, we process the matching islands of *T*_1_ and *T*_2_ by iterating over the good edges (of which there are *𝒪* (*n*)). We retrieve for a good edge the two islands to which it belongs in *T*_1_ (line 8) and in *T*_2_ (line 9). This is achieved in constant time using bipartition-to-island-pair mappings obtained during the tree traversal of *getIslands*() below.

For each of the matching island of *T*_1_ and *T*_2_ (line 10), the algorithm checks whether the pair has already been visited in a previous iteration of the loop (the same island pair can be visited from multiple good edges). If not, the current distance is updated by adding the number of bad edges in each island. Since these sizes are also pre-computed by *getIslands*(), this operation is in constant time as well.

The iteration over all good edges ends with lines 13-14, which account for a potentially required single substitution between corresponding islands, in case they have no label in common (i.e. they form a label-disjoint island pair). These operations can also be performed in constant time, giving an overall *𝒪* (*n*) runtime for the for-loop.

Finally, lines 16-19 are needed to handle the special case where there is no good edge between *T*_1_ and *T*_2_. In such a case, there is only one island per tree, which is matching.

### 4.2 The *getIsland*() function

We now detail the auxiliary function *getIslands*() (Algorithm 2). Recall that its goal is to identify the islands of an input tree, given a list of good edges. This is achieved through a single traversal of the tree in pre-order (we assume that the tree is arbitrarily rooted, and that the dummy root node has no label). In doing so, we identify the islands, which are separated by good edges, and keep track of (i) the set of labels found in each island (array *islLabels*); (ii) the number of bad edge in each island (array *islSizes*); (iii) the pair of islands associated with each bipartition (*bipart*2*isl*). These three data structures are initialised in lines 1-3. Note that the initial island is initialised to −1 because it will be incremented to 0 at the first step of the traversal.

Lines 4-20 define the recursive function used to traverse the tree. Because each good edge belongs to exactly two islands (Sect. 3), good edges can be used to identify the transition between two islands. By contrast, adjacent bad edges are by definition part of the same island. In our traversal, we thus check whether a particular node is attached to the previous island by a good edge (lines 7-15) or a bad edge (lines 16-20). If it is the former, we have just transitioned to a new island, and thus append new elements to the *islLabels* (line 8) and *islSizes* arrays (line 9).

Furthermore, we update the bipartition-to-island table with references to the two islands which are deliminated by the good edge. Since by definition a good edge induces the same bipartition in *T*_1_ and *T*_2_, the bipartition bitmask (a binary vector of the length *n* with 1 for all leaves present in the clade attached to the good edge) should either be the same for the good edge in *T*_1_ and *T*_2_, or bit-wise complementary if the rooting between *T*_1_ and *T*_2_ is on different sides of the good edge (as the tree data structure is arbitrarily rooted). We store the associated islands using both bitmasks, ensuring that the island which on same side as the root is listed first (lines 12-13).

If we encounter a node which is attached to the previous island by a bad edge, then it is still part of it, so we just update the set of leaf of the previous island (line 17), and increment its size counter by one (line 18).

All operations performed at each internal node are constant time, and the number of internal nodes is *𝒪* (*n*), so the time complexity of the tree traversal is done in time *𝒪* (*n*).

We provide an open source implementation of *LRF* in Python as part of the pyLabeledRF package (https://github.com/DessimozLab/pylabeledrf).

#### Algorithm 1 LRF(*T1; T2*)

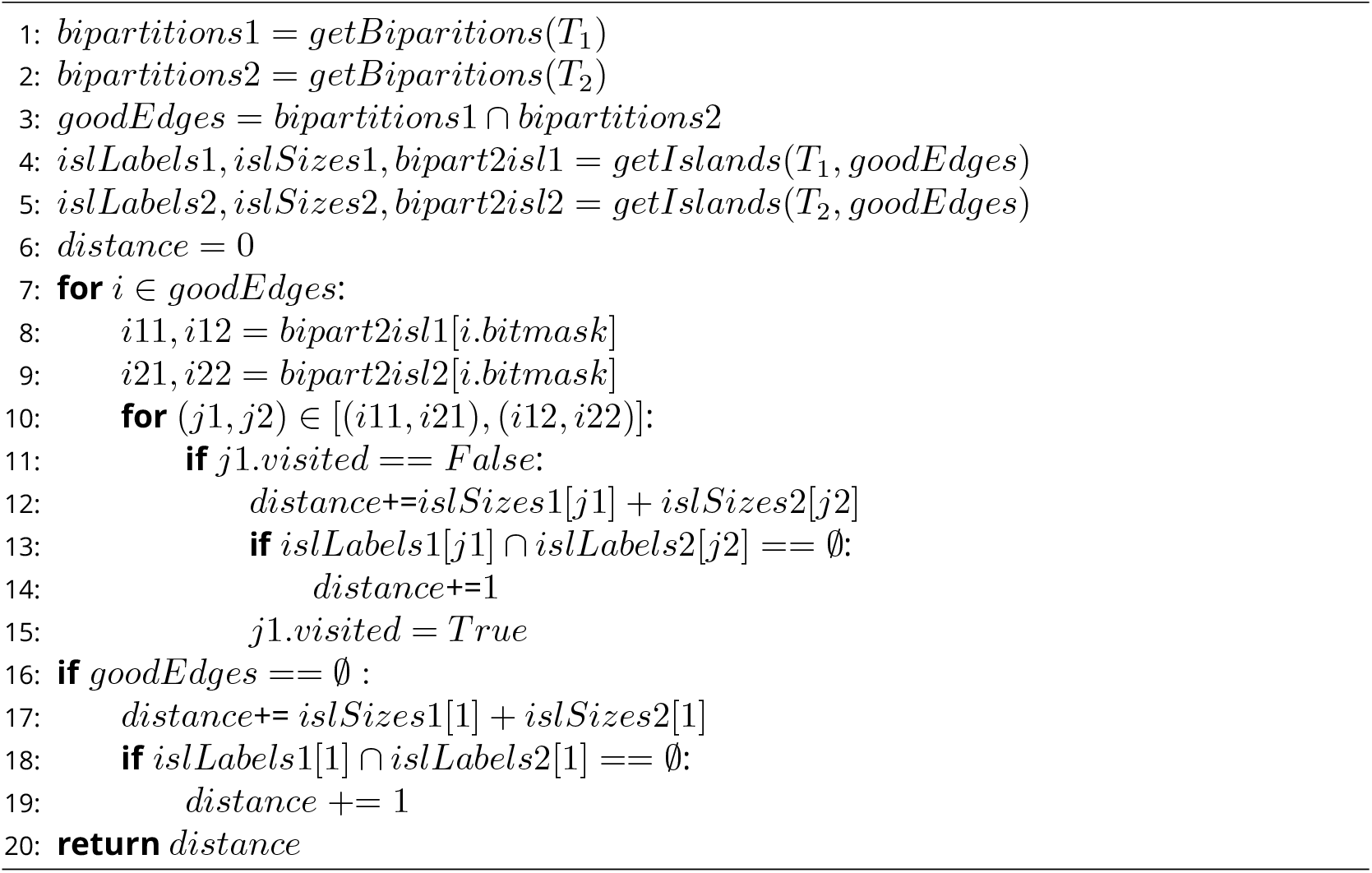

#### Algorithm 2 getIslands(*T; goodEdges*)

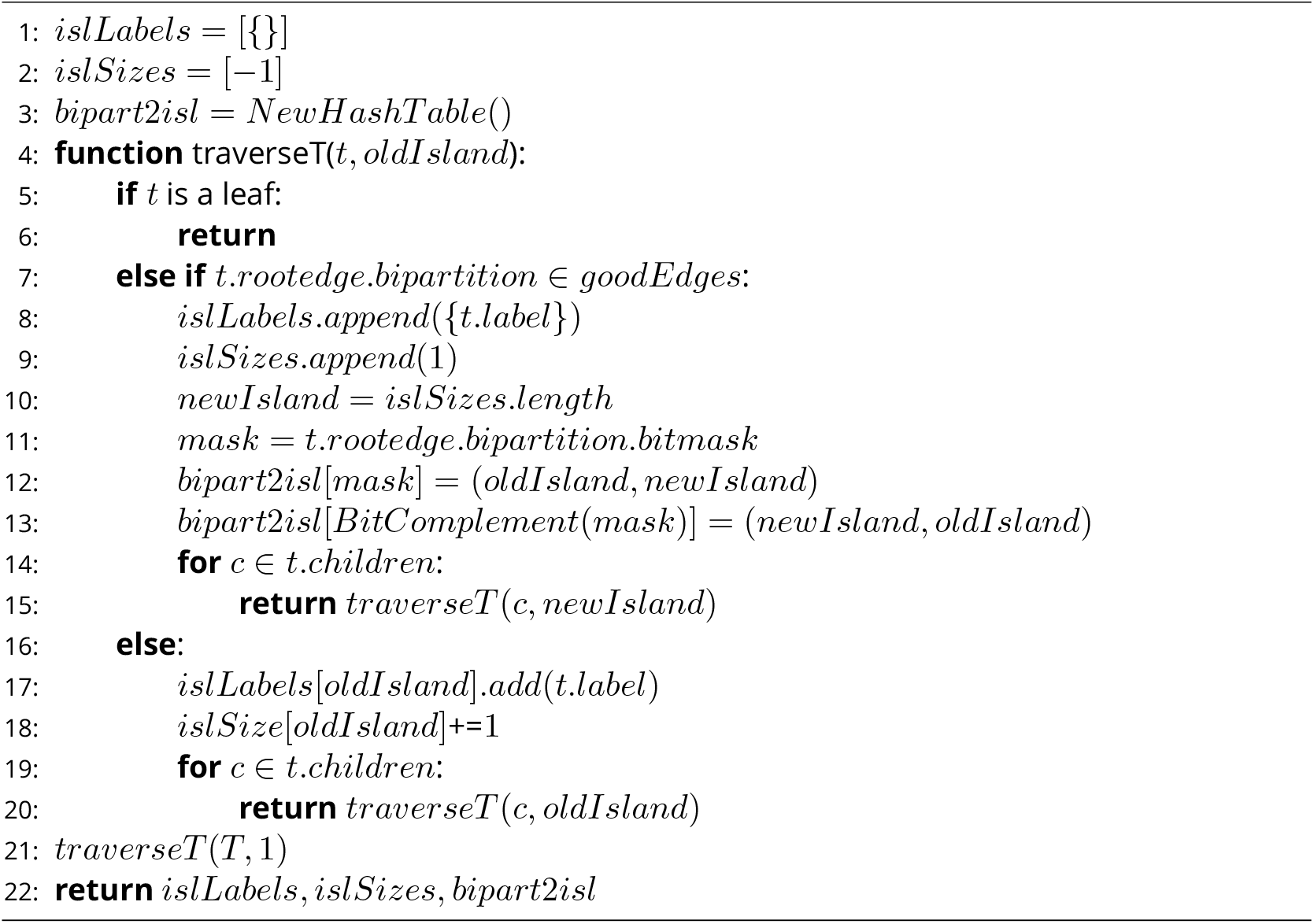

## 5 Experimental results

To illustrate the usefulness of *LRF*, we performed two experiments. First, we compared *LRF* with *RF* and *ELRF* on a labeled gene tree with random edits. Second, we used *LRF* to tackle an open question in orthology inference: does labeled gene tree inference benefit from denser taxon sampling?

### 5.1 Empirical comparison of *LRF* with *RF* and *ELRF*

To get a first sense of *LRF* ‘s ability to measure the actual number of edits between two trees, we performed a simulation study alongside *RF* and *ELRF*. We retrieved the labeled tree associated with human gene NOX4 from Ensembl release 99 (Yates et al., 2020), containing 182 genes, including speciation and duplication nodes. Next, we introduced a varying number of random edits, with 10 replicates, as follows: with probability 0.3, the label of one random internal node was substituted (from a speciation label into a duplication one or vice versa); the rest of the probability mass function was evenly distributed among all internal edges (each implying a potential node deletion) and all nodes of degree *>* 3 (each providing the opportunity of a potential node insertion). For *ELRF*, consistent with its underlying model, we added the requirement that edge removal only affect edges with adjacent nodes with the same label.

For each of *RF, LRF* and *ELRF*, we provide the distance as a function of the number of random edits (Fig. 6). As expected, the conventional *RF* distance returns the smallest values because it ignores labels; it however tracks quite well the expected number of node insertion and/or removal (dashed line). The two labeled *RF* alternatives performed similarly, but the heuristic for *ELRF* occasionally exceeded the true number of edit operations — a shortcoming that we do not have with *LRF*, as we have an exact algorithm for this distance. Both labeled *RF* variants tracked better the actual number of changes, until around 13 edits for *LRF* or *ELRF*, after which the minimum edit path starts to be often shorter than the actual sequence of random edits.

**Figure 6.**
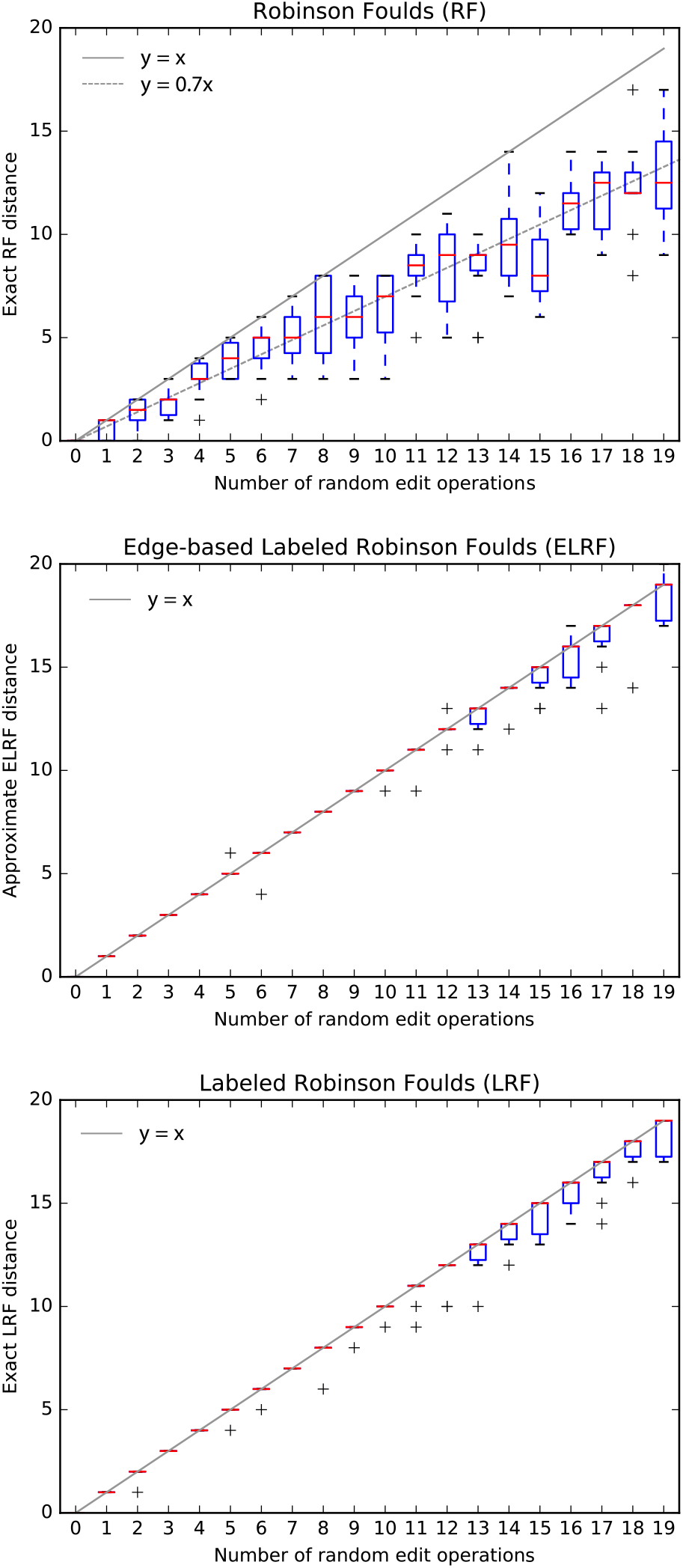
Empirical comparisons of the distance inferred for an increasing number of random edit operations (node insertion, deletion, substitution) on the NOX4 gene tree (182 leaves), using the classical RF distance (top), the ELRF approximation (Briand, Dessimoz, El-Mabrouk, Lafond, et al., 2020; middle), and the LRF exact distance (bottom).

### 5.2 The effect of denser taxon sampling on labeled gene tree inference

We used *LRF* to assess the effect of species sampling for the purpose of labeled gene tree reconstruction. Consider the problem of reconstructing a labeled tree corresponding to homologous genes from 10 species. Our question is: is it better to infer and label the tree using these 10 species alone, or is it better to use more species to infer and label the tree, and then prune the resulting tree to only contain the leaves corresponding to the original 10 species? While denser taxon sampling is known to improve unlabeled phylogenetic inference (Nabhan and Sarkar, 2011), we are not aware of any previous study on labeled gene tree inference.

First, using ALF (Dalquen et al., 2012), we simulated the evolution of the genomes of 100 extant species from a common ancestor genome containing 100 genes (*Parameters*: root genome with 100 genes of 432 nucleic acids each; species tree sampled from a birth-death model with default parameters; sequences evolved using the WAG model, with Zipfian gap distribution; duplication and loss events rate of 0.001). In the simulation, genes can mutate, be duplicated or lost. All the genes in the extant species can thus be traced back to one of these 100 ancestral genes and be assigned to the corresponding gene family. The 100 true gene trees, including speciation and duplication labels, are known from the simulation. However, in our run, one tree ended up containing only two genes (due to losses on early branches) and was thus excluded from the rest of the analysis.

To evaluate the inference process, among the 100 species, we randomly selected nested groups of 10, 20, 30, 40, 50, 60, 70, 80 and 90 species. We considered the 10 species in the first group as the species of interest. All other species were used to potentially improve the reconstruction of the gene trees for the first 10 genomes. Then, for each group, we aligned protein sequences translated from homologous genes using MAFFT L-INS-i (Katoh and Standley, 2013), inferred phylogenetic trees from the alignments using FastTree (Price et al., 2010), and annotated their nodes using the species overlap algorithm (Heijden et al., 2007) as implemented in the ETE3 python library (Huerta-Cepas et al., 2016). Finally, we pruned both the inferred gene trees and the true trees to include only proteins corresponding to the 10 species of interest.

We used *LRF* to assess the distance between the estimated and true labeled trees, for the various number of auxiliary genomes considered. For each scenario, we computed the mean *LRF* distance over all gene trees (Fig. 7). The mean error (expressed in *LRF* distance) decreases as the number of auxiliary species increases. This simple simulation study suggests that denser species sampling improves labeled gene tree inference.

**Figure 7.**
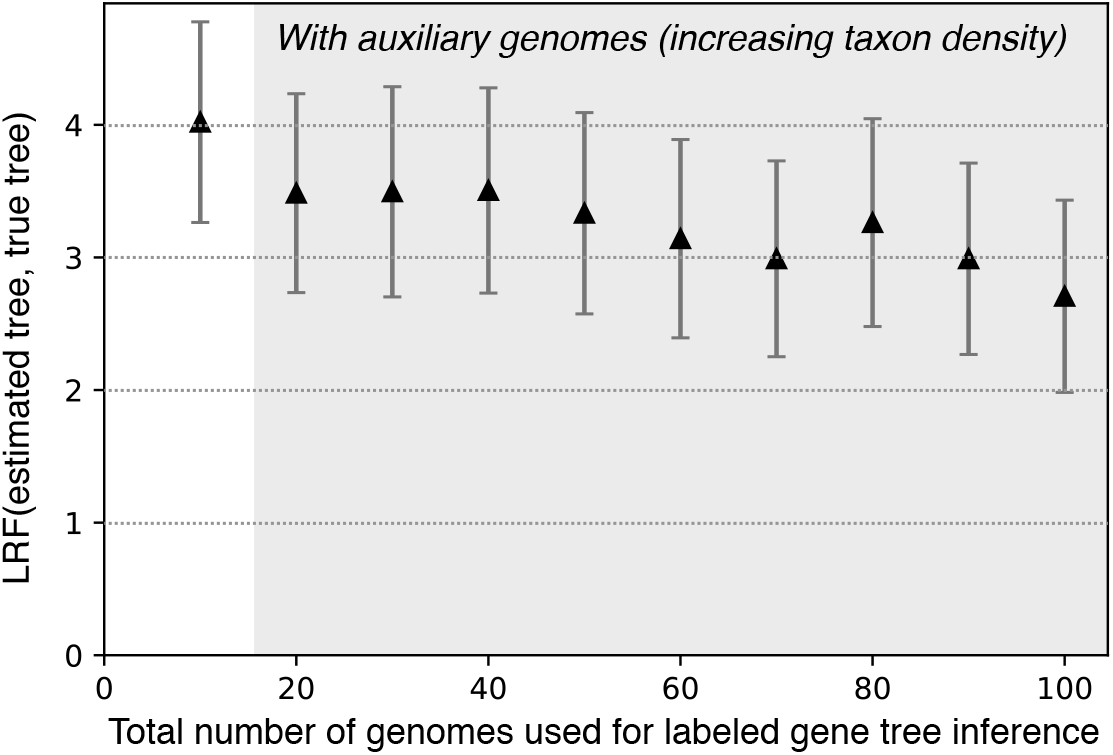
Denser taxon sampling decreases labeled tree estimation error: labeled gene trees reconstructed with an increasing number of auxiliary genomes (i.e. obtained by including the additional genomes during tree inference and labeling, followed by pruning) have a smaller *LRF* distance to the true trees. Error bars depict 95% confidence intervals around the mean.

## 6 Discussion and Conclusion

The *LRF* distance introduced here overcomes the major drawback of *ELRF*, namely the lack of an exact polynomial algorithm for the latter. Indeed, with *ELRF*, minimal edit paths can require contracting “good” edges, i.e., edges present in the two trees (Briand, Dessimoz, El-Mabrouk, Lafond, et al., 2020). By contrast, with *LRF*, we demonstrated that there is always a minimal path which does not contract good edges. Better yet, we proved that *LRF* can be computed exactly in linear time. The new formulation also maintains other desirable properties: being a metric, even for an arbitrary number of label types, and reducing to the conventional Robinson Foulds distance in the presence of trees with only one type of label.

Our experimental results provide a relationship between the number of random edits and the computed edit distances. At first sight, it may seem surprising that in a tree of 182 leaves, the minimum edit path under *LRF* or *ELRF* already starts underestimating the actual number of random edit operations after around 13 operations. However, this can be explained by the “birthday paradox” (Abramson and Moser, 1970): to be able to reconstruct the actual edit path, no two random edits should affect the same node. Yet the odds of having, among 13 random edits, at least two edits affecting the same internal node (among 179) is in fact substantial — approximately 36% in our case — just like the odds of having two people with the same birthday in a given group is higher than what most people intuit.

It has to be noted that *LRF* has the same limitations as *RF* regarding lack of robustness and skewed distribution. Moreover, like *RF* and *ELRF*, the main limitation of *LRF* is the lack of biological realism. For one thing, there is no justification to assign equal weight to the three kinds of edits in all circumstances. For instance, it is typically highly implausible to introduce a speciation node at the root of a subtree containing multiple copies of a gene in the same species. However, *LRF* complement analyses performed using more realistic models are either unavailable or too onerous to compute. In particular, the ability of *LRF* to support an arbitrary number of labels makes it applicable to gene trees containing more than just speciations and duplications, such as horizontal gene transfers or gene conversion events.

Finally, *LRF* constitutes a clear improvement over *RF* in the context of gene tree benchmarking, where trees inferred by various reconciliation models are compared using a distance measure (Altenhoff et al., 2016; Morel et al., 2019). Such an application was illustrated in the simulation study of the previous section, in which we observed that denser taxon sampling improved labeled tree inference computed using the widely used species overlap method. More work will be needed to assess the generality of this result.

## Data accessibility

Data and scripts for the experimental analyses are available online at https://doi.org/10.5281/zenodo.4329899

## Supplementary material

The software written in Python is available in the pylabeledrf repository at https://github.com/DessimozLab/pylabeledrf.

### Acknowledgements

We warmly thank Céline Scornavacca, Barbara Holland, Gabriel Cardona, Jean-Baka Domelevo Entfellner, and an anonymous reviewer for their detailed and constructive peer-reviews. The work was supported by Swiss National Science Foundation (SNSF) Professorship grant 183723 (to CD), by the Natural Sciences and Engineering Research Council of Canada (NSERC, to NEM), and Fonds de Recherche Nature et Technologies of Quebec (FRQNT, to NEM). Version 4 of this preprint has been peer-reviewed and recommended by Peer Community In Mathematical and Computational Biology (https://doi.org/10.24072/pci.mcb.100002).

## Conflict of interest disclosure

The authors of this preprint declare that they have no financial conflict of interest with the content of this article. Christophe Dessimoz serves on the PCI Math Comp Biol managing board and is a recommender, but was not involved in the handling of this manuscript.

## Notes

### Competing Interest Statement

The authors have declared no competing interest.

### Summary of Updates

Version 4 of this preprint has been peer-reviewed and recommended by Peer Community In Mathematical and Computational Biology (https://doi.org/10.24072/pci.mcb.100002)

## References

Abramson M and WOJ Moser (1970). More Birthday Surprises. The American mathematical monthly: the official journal of the Mathematical Association of America 77, 856–858. issn: 0002-9890, 1930-0972. doi: 10.2307/2317022.

Allen BL and M Steel (2001). Subtree transfer operations and their induced metrics on evolutionary trees. Annals of combinatorics 5, 1–15.

Altenhoff AM, B Boeckmann, S Capella-Gutierrez, DA Dalquen, T DeLuca, K Forslund, J Huerta-Cepas, B Linard, C Pereira, LP Pryszcz, et al. (2016). Standardized benchmarking in the quest for orthologs. Nature methods 13, 425–430.

Boussau B and C Scornavacca (2020). Reconciling gene trees with species trees. Phylogenetics in the Genomic Era, 3.2:1–3.2:23.

Briand S, C Dessimoz, N El-Mabrouk, M Lafond, and G Lobinska (2020). A Generalized Robinson-Foulds Distance for Labeled Trees. BMC Genomics 21. doi: 10.1186/s12864-020-07011-0.

Briand S, C Dessimoz, N El-Mabrouk, and Y Nevers (2020). A Linear Time Solution to the Labeled Robinson-Foulds Distance Problem. BioRxiv 2020.09.14.293522, ver. 4 peer-reviewed and recommended by PCI Mathematical & Computational Biology. doi: 10.1101/2020.09.14.293522.

Cardona G, M Llabrés, F Rosselló, and G Valiente (2010). Nodal distances for rooted phylogenetic trees. Journal of mathematical biology 61, 253–276.

Critchlow DE, DK Pearl, and C Qian (1996). The triples distance for rooted bifurcating phylogenetic trees. Systematic Biology 45, 323–334.

Dalquen DA, M Anisimova, GH Gonnet, and C Dessimoz (Apr. 2012). ALF–a simulation framework for genome evolution. Molecular biology and evolution 29, 1115–1123. issn: 0737-4038. doi: 10.1093/molbev/msr268.

Day WH (1985). Optimal algorithms for comparing trees with labeled leaves. Journal of classification 2, 7–28.

Estabrook GF, F McMorris, and CA Meacham (1985). Comparison of undirected phylogenetic trees based on subtrees of four evolutionary units. Systematic Zoology 34, 193–200.

Heijden RTJM van der, B Snel, V van Noort, and Ma Huynen (Jan. 2007). Orthology prediction at scalable resolution by phylogenetic tree analysis. BMC bioinformatics 8, 83. issn: 1471-2105. doi: 10.1186/1471-2105-8-83.

Hickey G, F Dehne, A Rau-Chaplin, and C Blouin (2008). SPR distance computation for un-rooted trees. Evolutionary Bioinformatics 4, EBO–S419.

Huerta-Cepas J, F Serra, and P Bork (June 2016). ETE 3: Reconstruction, Analysis, and Visualization of Phylogenomic Data. en. Molecular biology and evolution 33, 1635–1638. issn: 0737-4038, 1537-1719. doi: 10.1093/molbev/msw046.

Jiang BDXHT, M Li, J Tromp, and L Zhang (2000). On computing the nearest neighbor interchange distance. In: Discrete Mathematical Problems with Medical Applications: DIMACS Workshop Discrete Mathematical Problems with Medical Applications, December 8-10, 1999, DIMACS Center. Vol. 55. American Mathematical Soc., p. 125.

Katoh K and DM Standley (Apr. 2013). MAFFT multiple sequence alignment software version 7: improvements in performance and usability. Molecular biology and evolution 30, 772–780. issn: 0737-4038, 1537-1719. doi: 10.1093/molbev/mst010.

Lin Y, V Rajan, and BM Moret (2012). A metric for phylogenetic trees based on matching. IEEE/ACM Transactions on Computational Biology and Bioinformatics (TCBB) 9, 1014–1022.

Mittal S and G Munjal (2015). Tree Mining and Tree Validation Metrics: A Review. IOSR: Journal of Computer Engineering, 31–36.

Moon J and O Eulenstein (2018). Cluster matching distance for rooted phylogenetic trees. In: International Symposium on Bioinformatics Research and Applications. Springer, pp. 321–332.

Morel B, AM Kozlov, A Stamatakis, and GJ Szöllősi (2019). GeneRax: A tool for species tree-aware maximum likelihood based gene tree inference under gene duplication, transfer, and loss. bioRxiv. doi: 10.1101/779066. eprint: https://www.biorxiv.org/content/early/2019/09/26/779066.full.pdf.

Nabhan AR and IN Sarkar (Mar. 2011). The impact of taxon sampling on phylogenetic inference: a review of two decades of controversy. Briefings in bioinformatics. issn: 1467-5463. doi: 10.1093/bib/bbr014.

Pattengale ND, EJ Gottlieb, and BM Moret (2007). Efficiently computing the Robinson-Foulds metric. Journal of Computational Biology 14, 724–735.

Price MN, PS Dehal, and AP Arkin (2010). FastTree 2–approximately maximum-likelihood trees for large alignments. PloS one 5, e9490.

Robinson DF and LR Foulds (1981). Comparison of phylogenetic trees. Mathematical biosciences 53, 131–147.

Schwarz S, M Pawlik, and N Augsten (2017). A new perspective on the tree edit distance. In: International Conference on Similarity Search and Applications. Springer, pp. 156–170.

Vilella A, J Severin, A Ureta-Vidal, L Heng, R Durbin, and E Birney (2009). EnsemblCompara gene trees: Complete, duplication-aware phylogenetic trees in vertebrates. Genome Research 19, 327–335.

Yates AD, P Achuthan, W Akanni, J Allen, J Allen, J Alvarez-Jarreta, MR Amode, IM Armean, AG Azov, R Bennett, et al. (2020). Ensembl 2020. Nucleic acids research 48, D682–D688.

Zhang K (1993). A new editing based distance between unordered labeled trees. In: Annual Symposium on Combinatorial Pattern Matching. Springer, pp. 254–265.

Zhang K (1996). A constrained edit distance between unordered labeled trees. Algorithmica 15, 205–222.

Zhang K and D Shasha (1989). Simple fast algorithms for the editing distance between trees and related problems. SIAM journal on computing 18, 1245–1262.

Zhang K, R Statman, and D Shasha (1992). On the editing distance between unordered labeled trees. Information processing letters 42, 133–139.

